# Teneurins Are SPARCL1 Receptors

**DOI:** 10.64898/2026.07.13.738299

**Authors:** Xuchen Zhang, Xudong Chen, Yi Miao, Thomas C. Südhof

## Abstract

Extensive experiments document that SPARCL1, a secreted protein that is produced primarily by astrocytes in brain and endothelia throughout the body and that is also known as Hevin, enhances synapse formation. However, the mode of action of SPARCL1 at synapses remains unclear owing to divergent results in the literature. Here, we use cultured neurons from newborn male and female mouse embryos to show that the C-terminal follistatin-like and Ca^2+^-binding domains of SPARCL1, which account for only 35% of the total SPARCL1 sequence, are sufficient to potently enhance synapse numbers. SPARCL1 acts at nanomolar concentrations at which SPARCL1 does not robustly bind to neurexins, neuroligins or neurexin/neuroligin complexes but avidly interacts with all teneurins. Strikingly, the follistatin-like domain of SPARCL1 on its own strongly binds to teneurins but is unable to stimulate synapse formation. Only when combined with the SPARCL1 Ca^2+^- binding domain does the follistatin-like domain induce synapses, suggesting that SPARCL1 enhances synapse numbers by binding to teneurins via its C-terminal follistatin-like domain and by activating synapse formation via its Ca^2+^-binding domain.

**SIGNIFICANCE STATEMENT:** SPARCL1 (also known as Hevin) is a synaptogenic factor that is produced primarily by astrocytes in brain, and that enhances synapse formation. How SPARCL1 acts at synapses, however, remains unclear because divergent results describe its binding partners at synapses and the sequences involved in its synaptogenic activity remain unclear. In the present study, we show that SPARCL1 avidly binds to the presynaptic teneurins adhesion molecules, that this binding is mediated by its small follistatin-like domain, and that its synaptogenic activity requires both its follistatin-like and its Ca^2+^-binding EC domains. Thus, our results suggest that SPARCL1 is recruited to developing synapses by binding of its follistatin-like domain to teneurins and then induces synapse assembly via its Ca^2+^-binding domain.

## INTRODUCTION

Synapses are induced and maintained in brain at least in part by astrocyte-derived synaptogenic factors. (Chung WS et al., 2024;Fossati G et al., 2020;Khaspekov LG and Frumkina LE, 2023). Pioneering work by Ben Barres’ laboratory identified SPARCL1 (a.k.a. Hevin) as a major synaptogenic factor from astrocytes (Kucukdereli H et al., 2011). SPARCL1 is composed of an N-terminal signal peptide, a largely natively unfolded N-terminal sequence (approximately 420 residues), and a highly conserved C-terminal sequence (approximately 230 residues) comprising a follistatin-like (FS domain) and a Ca^2+^- binding domain (EC domain) (Girard JP and Springer TA, 1995) (Fig. 1A). The FS and EC domains of SPARCL1 are highly homologous to those of SPARC and of testicans (Bradshaw AD, 2012). SPARCL1 is primarily expressed by astrocytes but also abundantly produced by vascular cells in and outside of brain (Girard JP and Springer TA, 1996).

**Figure 1.**
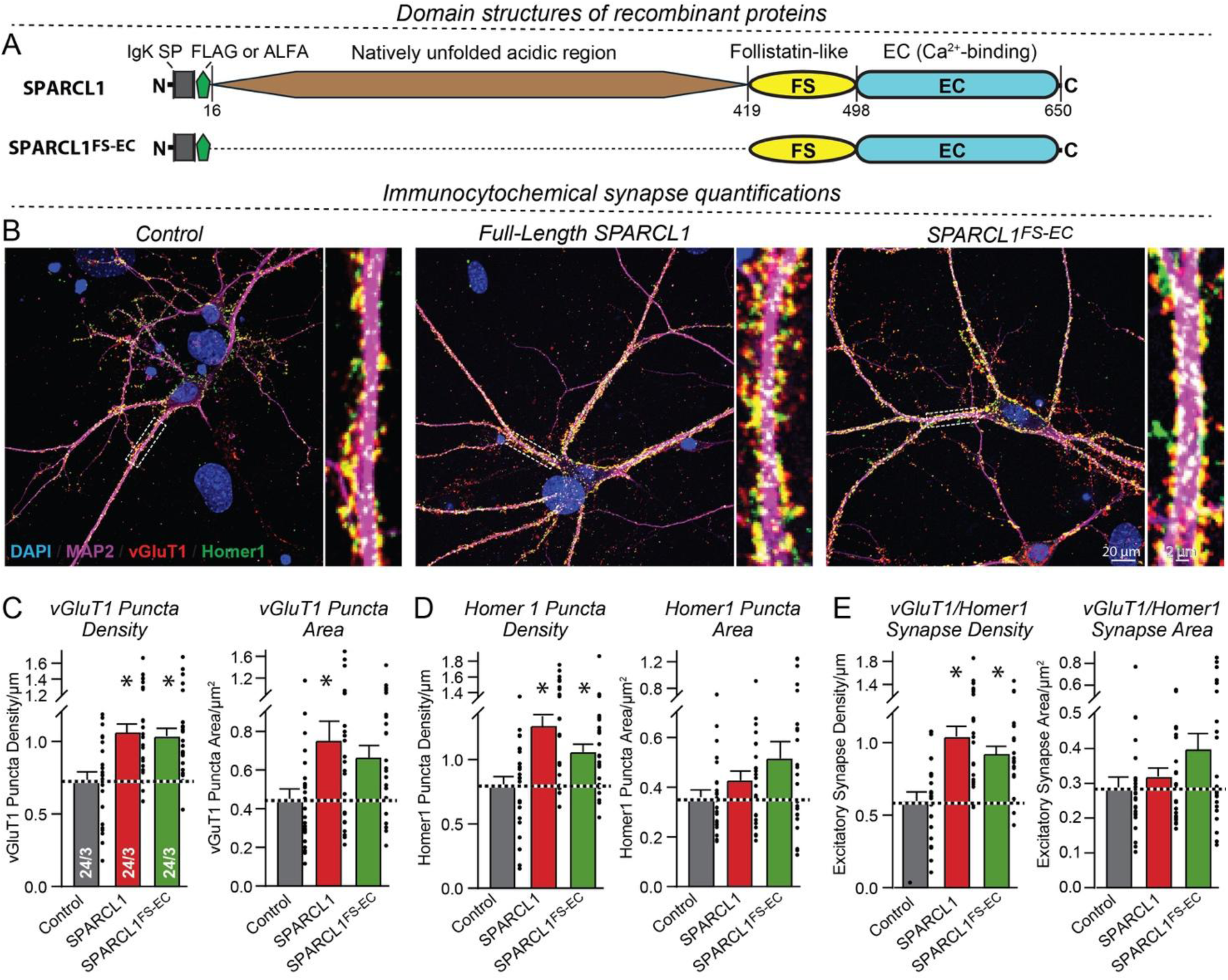
A SPARCL1 fragment comprising only its C-terminal follistatin-like (FS) and Ca^2+^-binding (EC) domains potently increases synapse numbers in cultured neurons. (A) Domain structures of full-length SPARCL1 and of its C-terminal fragment (SPARCL1^FS-EC^) composed of its follistatin-like (FS) and Ca^2+^-binding (EC) domains. (B) Representative confocal images of mixed neuron-glia cultures treated with control medium or medium containing 50 nM recombinant full-length FLAG-tagged SPARCL1 or SPARCL1^FS-EC^. Treatments of hippocampal cultures from newborn mice started at DIV6 or DIV7, media were refreshed at DIV10 or DIV11, and neurons were analyzed by immunocytochemistry for vGluT1 (red), Homer1 (green), and MAP2 (magenta), and additionally stained with DAPI (blue) at DIV14 or DIV15. For each condition, overviews are shown on the left and higher-magnification images taken from the boxed areas on the right. (**C-E**) Summary graphs of the synapse density and size of excitatory synapses show that both full- length SPARCL1 and SPARCL1^FS-EC^ robustly increased the excitatory synapse density and modestly increased the apparent synapse size in the experiments described in B (**C** & **D**, quantifications of the vGluT1- and Homer1-positive puncta, respectively; **E**, quantifications of synaptic puncta that are positive for both vGluT1 and Homer1, with all images recorded with identical parameters). Data are means ± SEM (numbers of cells and experiments are indicated in bars); *P < 0.05 [one-way ANOVA with post-hoc Tukey tests]. For single channel images, see Figure S1.

SPARCL1 was initially localized to synapses (Lively S and Brown IR, 2008;Lively S et al., 2007). Using cultured retinal ganglion cells, Kucukdereli et al. (2011) then showed that full-length SPARCL1 and a central SPARCL1 sequence potently induced synapse formation, whereas a SPARCL1 fragment comprising its C-terminal FS and EC domains antagonized synapse formation (Kucukdereli H et al., 2011). These results were expanded by Singh et al. (2016) who reported that cleaving SPARCL1 at residue 350 (in the central SPARCL1 sequence 80 residues N-terminal to the FS domain), blocks SPARCL1’s synaptogenic activity. Truncation of SPARCL1 at residue 459 in the FS domain, conversely, did not abolish synaptogenic activity (Singh SK et al., 2016). Moreover, Singh et al. (2016) found that a SPARCL1 fragment N-terminal to the FS domain directly binds to neurexin-1α (Nrxn1α) and to neuroligin-1B (Nlgn1B), which are pre- and postsynaptic adhesion molecules, respectively. Via this binding, SPARCL1 was proposed to form a trimeric complex, suggesting that a synaptogenic SPARCL1 sequence in the region N-terminal to the FS domain acts in synaptogenesis by strengthening the Nrxn1α-Nlgn1B complex and that this action is antagonized by the FS and EC domains of SPARCL1.

Recent studies confirmed a robust synaptogenic activity of SPARCL1 (Gan KJ and Sudhof TC, 2019;Gan KJ and Sudhof TC, 2020). Puzzlingly, however, the genetic deletion of all neurexins or all neuroligins did not significantly impair the synaptogenic action of SPARCL1, suggesting that SPARCL1 does not require neurexins or neuroligins for its synaptogenic activity (Gan KJ and Sudhof TC, 2020). To complicate matters, a subsequent biophysical paper confirmed SPARCL1 binding to Nrxn1a, Nlgn1, and Nlgn2 but localized the Nrxn1a- and Nlgn1-binding site of SPARCL1 to its C-terminal fragment containing the FS and EC domains instead of the sequence N-terminal to the FS domain (Fan S et al., 2021).

Thus, the present data provide incompatible results about SPARCL1’s synaptogenic activity in that different papers identify distinct Nrxn1α- and Nlgn1B-binding sequences in SPARCL1 (Fan S,Gangwar SP,Machius M and Rudenko G, 2021) and yet another paper shows that the neurexin and neuroligin deletions have no effect on the synaptogenic activity of SPARCL1 (Gan and Sudhof, 2020). Moreover, two papers Kucukdereli et al. (2011) and Singh et al. (2016) reported that the C-terminal SPARCL1 fragment inhibits the synaptogenic activity of SPARCL1, whereas another study Fan et al. (2021) revealed that the FS and EC domain fragment of SPARCL1 mediates binding to Nrxn1a and Nlgn1B that supposedly induces synaptogenesis. In addition, neurexins and neuroligins form high-affinity complexes (Comoletti D et al., 2006), raising the question of how low-affinity binding by SPARCL1 to neurexins and neuroligins could strengthen their high-affinity complexes.

Here, we have aimed to examine these questions. We find that SPARCL1 potently induces synapse formation via its FS and EC domains without its N-terminal sequences. Moreover, we unexpectedly identified strong binding of SPARCL1 to teneurins, which are presynaptic adhesion molecules (Zhang X et al., 2025; Zhang X et al., 2022), instead of neurexins and neuroligins. Finally, we found that the small SPARCL1 FS domain mediates teneurins binding but requires the EC domain for synaptogenicity. Our data suggest a model whereby the conserved SPARCL1 FS and EC domains induce synapse formation by coupling FS domain-mediated teneurins binding to the synaptogenic activity of the EC domain.

## RESULTS

### The SPARCL1 FS and EC domains drive synapse formation

As a first goal, we aimed to determine which domains of SPARCL1 stimulate synapse assembly. We hypothesized that the conserved domains of SPARCL1 which are present in all SPARC protein family members, namely its follistatin-like (FS) and Ca^2+^-binding (EC) domains (Fig. 1A), might be sufficient because its longer N-terminal sequence appears to be natively unfolded and less conserved evolutionarily. Thus, we generated recombinant proteins composed of full-length SPARCL1 and of the FS and EC domain fragment of SPARCL1 (called SPARCL1^FS-EC^; Fig. 1A).

We generated primary mixed hippocampal neuron-glia cultures from newborn mice, treated the cultures with a control medium or with 50 nM SPARCL1 or SPARCL1^FS-EC^ from DIV6 or DIV7, refreshed the medium at DIV10 or DIV11, and analyzed the cultures at DIV14 or DIV15 by immunocytochemistry for the presynaptic marker vGluT1, the postsynaptic marker Homer1, and the dendritic marker MAP2 (Fig. 1B, Fig. S1).

Both full-length SPARCL1 and SPARCL1^FS-EC^ increased the synapse density approximately 40-50% and elevated the apparent synapse size by a slightly lower amount (Fig. 1B-E). Here, the synapse density was measured either separately as the number of vGluT1 or Homer1 immunoreactive puncta per dendrite length (Fig. 1B-D) or as the number of puncta that are immunoreactive for both vGluT1 and Homer1 (Fig. 1E), with similar results for all three approaches.

The finding that the SPARCL1^FS-EC^ fragment is sufficient to promote synapse formation differs from a previous study which localized the synaptogenic activity of SPARCL1 to a sequence N-terminal of the FS domain (Singh et al., 2016). To ensure the reliability of our findings beyond using a rigorous approach to quantifying the synapse density (Fig. 1), we therefore analyzed the effect of full-length SPARCL1 or of SPARCL1^FS-EC^ on the network activity of neurons in neuron-glia cultures as a proxy of synaptic function (Fig. 2). We employed Ca^2+^-imaging of neurons using genetically encoded gCamp-6m driven by the human synapsin-1 promoter (Sudhof TC, 1990). With this approach, we can reliably measure the firing of neurons in the culture (Fig. 2A, B) and analyze parameters such as the frequency and amplitude of spikes and their synchronicity in the network (Fig. 2C, D).

**Figure 2.**
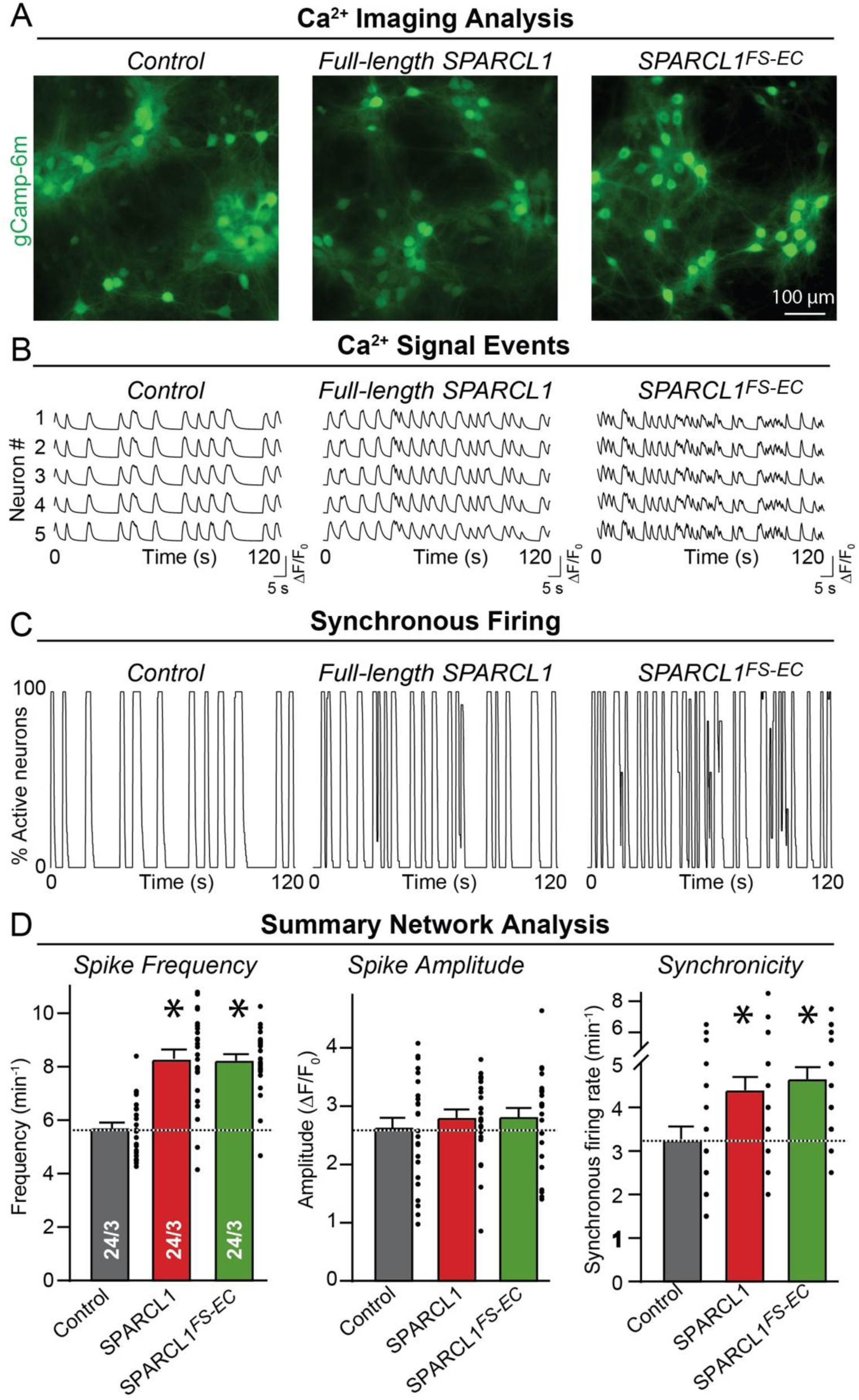
The SPARCL1 fragment composed of only its C-terminal follistatin-like and Ca^2+^-binding domains robustly enhances neuronal network activity as a proxy for synaptic transmission. (A) Representative images from Ca^2+^-monitoring experiments in mixed hippocampal neuron-glia cultures expressing GCaMP6m (green) under control of the human Synapsin-1 promoter (Sudhof TC, 1990). Cultures were treated with control or SPARCL1- or SPARCL1^FS-EC^-containing media as described for Figure 1B, except that the cultures were infected at DIV4 with lentiviruses expressing GCaMP6m, and that the SPARCL1 used was ALFA-tagged instead of FLAG-tagged. (B) Representative traces illustrating the extraction of GCaMP6m signals from raw Ca^2+^-imaging recordings of individual neurons. (C) Representative analyses of the synchronous firing rate of neurons in a culture computed from raw Ca^2+^-imaging recordings. (D) Quantifications of the spike frequency representing action potentials (left), amplitude (middle), and synchronous firing rate (right) of the network activity monitored by Ca^2+^-imaging show that full- length SPARCL1 and the SPARCL1 ^FS-EC^ fragment increase the frequency and synchronicity of neuronal firing in hippocampal cultures. Data are means ± SEM (numbers of cells and experiments are indicated in bars), with *P < 0.05 [one-way ANOVA with post-hoc Tukey tests].

Our results show that full-length SPARCL1 and the SPARCL1^FS-EC^ fragment have the same effect on neural network activity in the cultures, increasing the frequency and synchronicity of firing approximately 30-40% (Fig. 2D). These results confirm that the SPARCL1 or SPARCL1^FS-EC^ fragment is equally active in promoting synapse formation as full-length SPARCL1, consistent with the high degree of conservation of the FS and EC domains of SPARCL1.

### The SPARCL1 FS and EC domains bind to teneurins

Based on previous results suggesting that SPARCL1 binds to both neurexins and neuroligins (Singh SK et al., 2016; Fan S et al., 2021), we hypothesized that the SPARCL1^FS-EC^ fragment may drive synapse formation by binding to neurexins and neuroligins. Note, however, that only Fan et al. (2021) localized the neurexin- and neuroligin-binding site of SPARCL1 to the SPARCL1^FS-EC^ fragment, whereas Singh et al. (2016) localized the neurexin- and neuroligin-binding activities of SPARCL1 to a more N-terminal sequence.

To pursue the neurexin- and neuroligin-binding hypothesis, we first examined the binding of SPARCL1to two major splice variants (SS4- and SS4+) of neurexin-1α to -3α and of neurexin-1β to -3β as well as to neuroligin-1 to -3 in the context of a cellular environment. We co-expressed the 12 different neurexins and three neuroligins in HEK293 cells together with tdTomato, reacted the cells with 50 nM SPARCL1 in the medium overnight, fixed the cells, and visualized the surface neurexins and neuroligins as well as the bound SPARCL1 by immunocytochemistry (Fig. 3). As a supposed negative control, we used teneurin-3 (Tenm3).

**Figure 3.**
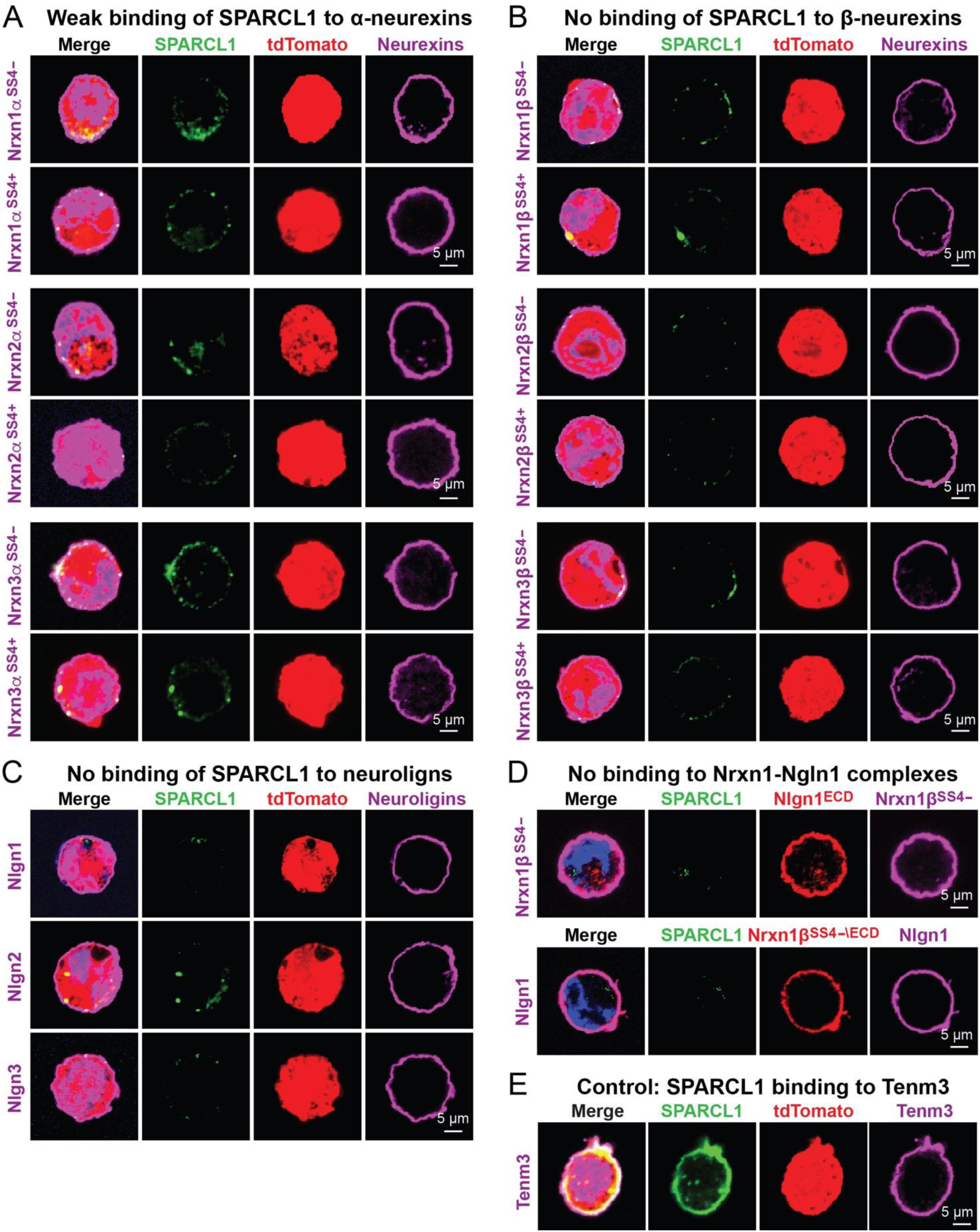
Cell-surface binding assays fail to detect robust SPARCL1 binding to neurexins, neuroligins, or the neurexin-1/neuroligin-1 complex. (**A**,**B**) Cell-surface binding assays testing the recruitment of recombinant full-length ALFA-tagged SPARCL1 (50 nM) to various isoforms of HA-tagged α-neurexins (A) or V5-tagged β-neurexins (B) that are expressed on the surface of HEK293T cells show that SPARCL1 does not robustly bind to α- or β-neurexins. HEK293T cells were co-transfected with the indicated α- and β-neurexins and tdTomato, incubated for 24 hours at 37 ^0^C with SPARCL1, and analyzed by staining the cell surfaces for SPARCL1 (green) and α- and β-neurexins (magenta). Representative images are from experiments independently repeated at least 3 times. (C) Cell-surface binding assays as described for A & B but analyzing HA-tagged neuroligins instead of neurexins do not reveal any binding of SPARCL1 to neuroligin-1 to -3 (Nlgn1-3). Representative images are from experiments independently repeated at least 3 times. (D) Cell-surface binding assays as described for A-C but analyzing surface-exposed Nrxn1β/Nlgn1 complexes without tdTomato co-expression instead of examining neurexins or neuroligins separately fail to detect SPARCL1 binding to surface neurexin/neuroligin complexes. HEK293T cells expressing either V5-tagged Nrxn1β (top row) or Nlgn1 (bottom row) were incubated for 24 hours at 37 ^0^C with the recombinant extracellular domains of Nlgn1 or Nrxn1β, respectively (both at 50 nM), to form the neurexin/neuroligin complexes on the cell surface. Cells were then washed, incubated with purified SPARCL1, and stained by immunocytochemistry for SPARCL1, Nlgn1 and Nrxn1β, with the co-labeling for Nlgn1 and Nrxn1β demonstrating that the surface neurexin-neuroligin complex had been formed but no significant SPARCL1 binding was detected. Representative images are from experiments independently repeated at least 3 times. (E) Cell-surface binding assays performed as described for A & B but using HEK293T cells expressing teneurin-3 (Tenm3) reveal strong binding of SPARCL1 to Tenm3. Representative images are from experiments independently repeated at least 3 times.

Strikingly, we detected no robust binding of SPARCL1 to any neurexin isoform or any neuroligin. For alpha-neurexins, we observed possibly weak binding (Fig. 3A), whereas unexpectedly, we found strong binding of SPARCL1 to Tenm3, the supposed negative control (Fig. 3E, Fig. S2). Note that these experiments were performed in a cellular environment at a very low SPARCL1 concentration (50 nM), identical to the concentration used for the synaptogenesis assays in Figures 1 and 2, which may explain why we were unable to reproduce previous studies (Fan S,Gangwar SP,Machius M and Rudenko G, 2021).

Given the unexpected binding of SPARCL1 to Tenm3, we next asked whether SPARCL1 might bind to all teneurin isoforms (Tenm1 to Tenm4). Here, we now used SPARCL1 binding to neurexin-1a (Nrxn1a) and cerebellin-1 (Cbln1) binding to teneurins as negative controls, and controlled the negative controls in turn by confirming Cbln1 binding to Nrxn1a (Fig. 4). We observed robust binding of SPARCL1 to all teneurins but not to Nrxn1a (Fig. 4A, B), whereas Cbln1 did not bind to any teneurin but robustly bound to Nrxn1a (Fig. 4C, D).

**Figure 4.**
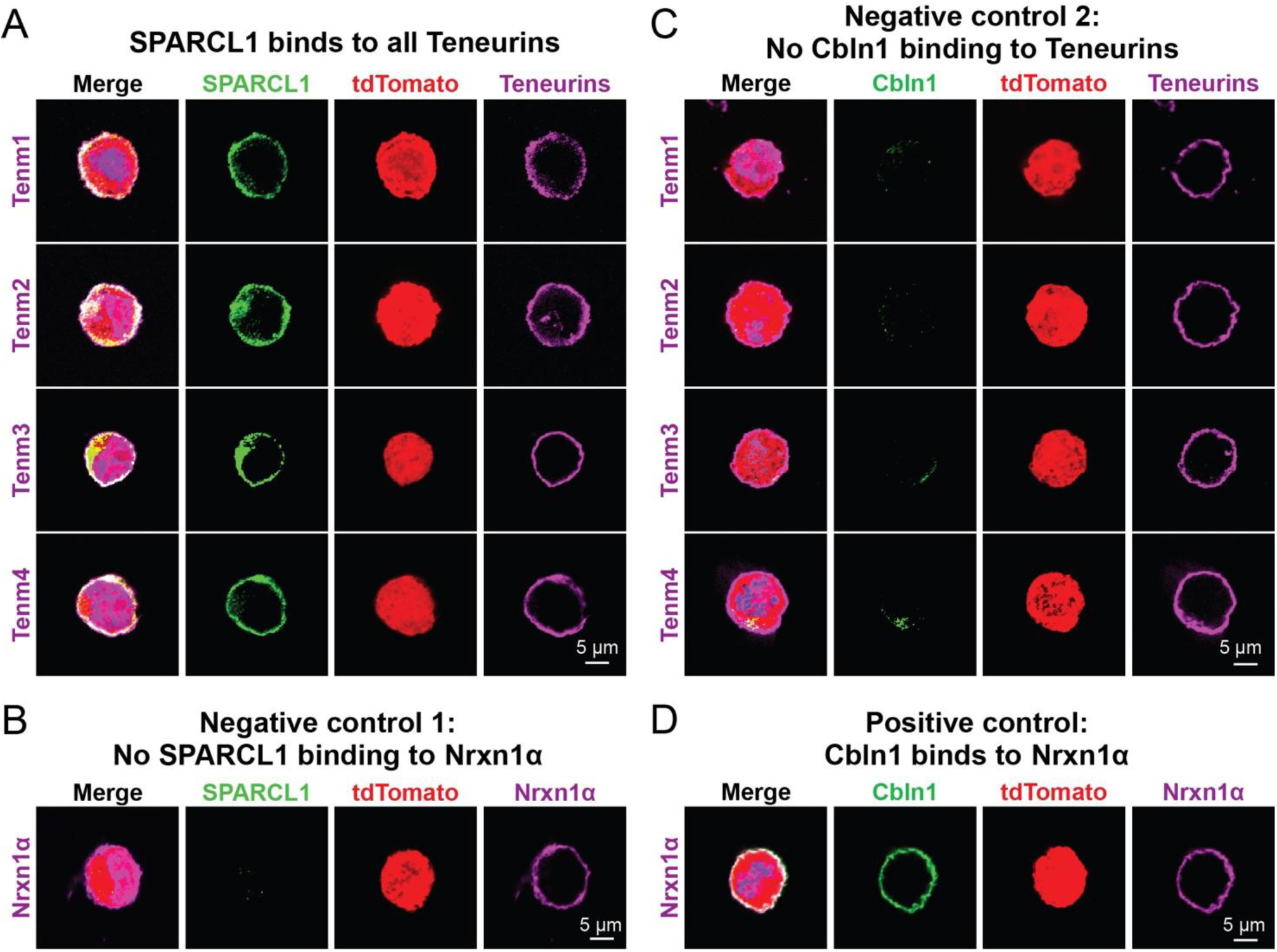
SPARCL1 robustly binds to all teneurins. (A) Cell-surface binding assays, performed as described for Figure 3A & B but analyzing teneurins instead of neurexins, reveal robust recruitment of SPARCL1 (50 nM) to full-length Tenm1-4 expressed on the surface of transfected HEK293T cells. HA-tagged teneurins and bound FLAG-tagged SPARCL1 were visualized by immunocytochemistry. (**B**-**D**) Negative and positive controls for SPARCL1 binding to teneurins using the same surface-binding assay as in panel A but demonstrating that neurexin1α (Nrxn1α) shows no recruitment of SPARCL1 (B) and that teneurins cannot bind FLAG-tagged Cerebellin-1 (Cbln1) (C) but that Cbln1 avidly binds to neurexin-1α (Nrxn1α) (D).

To confirm SPARCL1 binding to teneurins and explore whether the SPARCL1^FS-EC^ fragment may also bind to teneurin, we measured the binding of purified recombinant full-length SPARCL1 and of the SPARCL1^FS-EC^ fragment as a function of concentration to Tenm1 expressed on the surface of a HEK293 cell (Fig. 5). Both full-length SPARCL1 and of the SPARCL1^FS-EC^ fragment bound with similar affinities (∼73 nM and ∼88 nM, respectively), suggesting that the observed teneurin binding by SPARCL1 is mediated by its two C-terminal domains (Fig. 5).

**Figure 5.**
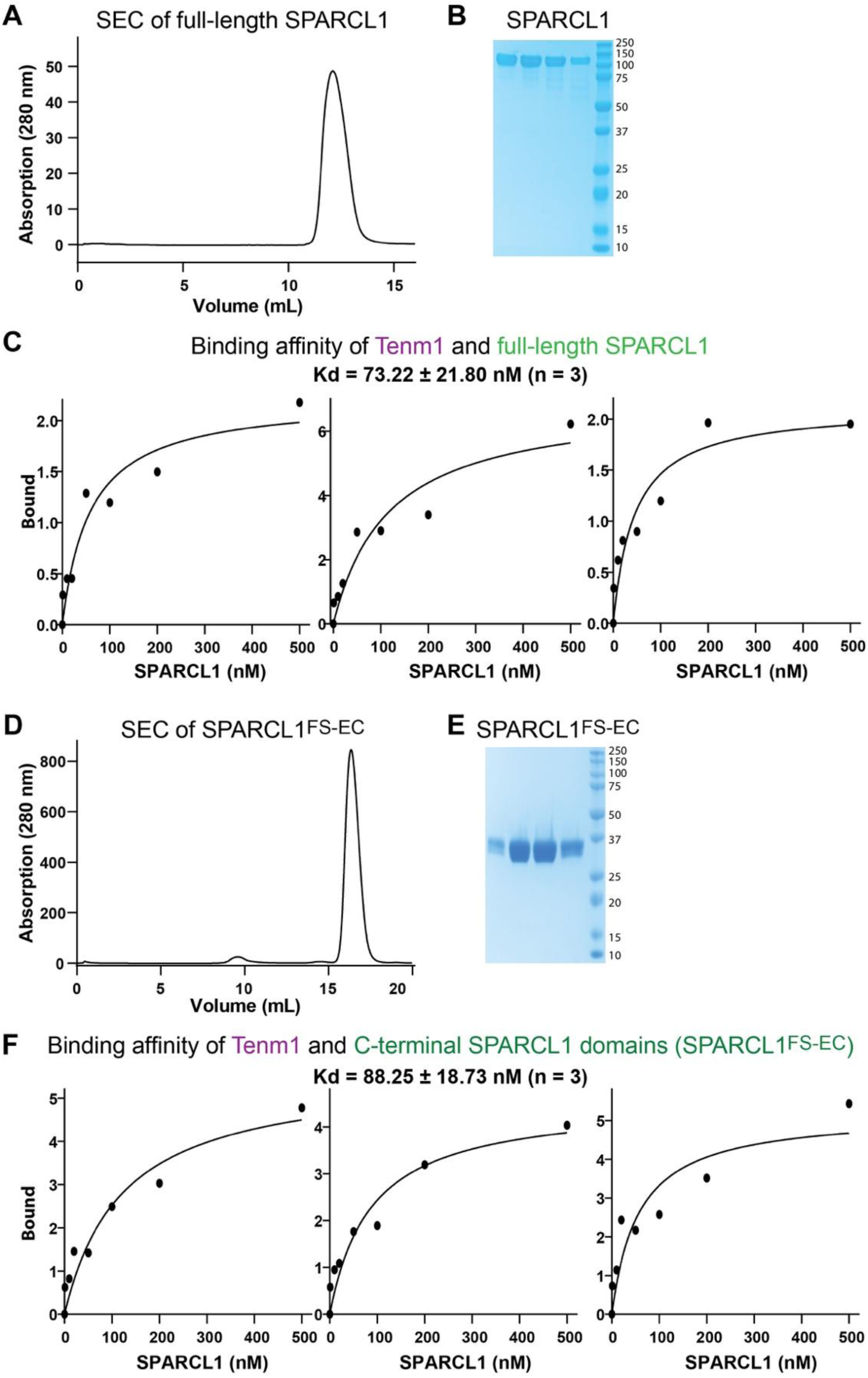
Full-length SPARCL1 and the SPARCL1^FS-EC^ fragment bind to teneurin-1 (Tenm1) on the cell surface with nanomolar affinities. (**A**,**B**) Purification of recombinant full-length FLAG-tagged SPARCL1 by size-exclusion chromatography (SEC) (A) and analysis of the purified protein by SDS-PAGE and Coomassie Blue staining (B). (**C**) Three independent experiments analyzing the binding of full-length SPARCL1 to teneurin-1 (Tenm1) expressed on the surface of transfected HEK293T cells. Different concentrations (1 nM, 10 nM, 20 nM, 50 nM, 100 nM, 200 nM, 500 nM) of full-length SPARCL1 were incubated with HEK293T cells transfected with Tenm1 or CMV vector alone (control), and the relative amounts of bound SPARCL1 was measured using immunocytochemistry. The control signal observed in CMV vector- transfected HEK293T cells was subtracted to calculate net binding signals. The calculated K_d_ was determined by three independent experiments. (**D**-**F**) The same experiments as shown for A-D, except that the C-terminal FLAG-tagged SPARCL1^FS-EC^ fragment was investigated instead of full-length SPARCL1.

### The SPARCL1 FS domain alone binds to teneurins but requires the EC domain for synaptogenic activity

We next asked whether teneurin-binding by the SPARCL1^FS-EC^ fragment is mediated by the FS or EC domain. To address this question, we produced each domain separately as a recombinant protein (Fig. 6A-C). We then tested each protein for binding to all teneurins expressed on the surface of HEK293 cells, using Nrxn1a as a negative control (Fig. 6D, E). Strikingly, the small FS domain (80 residues) was sufficient for binding to all teneurins at a concentration of 50 nM, whereas no binding of the EC domain to teneurins was detected (Fig. 6D, E). Moreover, no binding of either domain to Nrxn1a was found.

**Figure 6.**
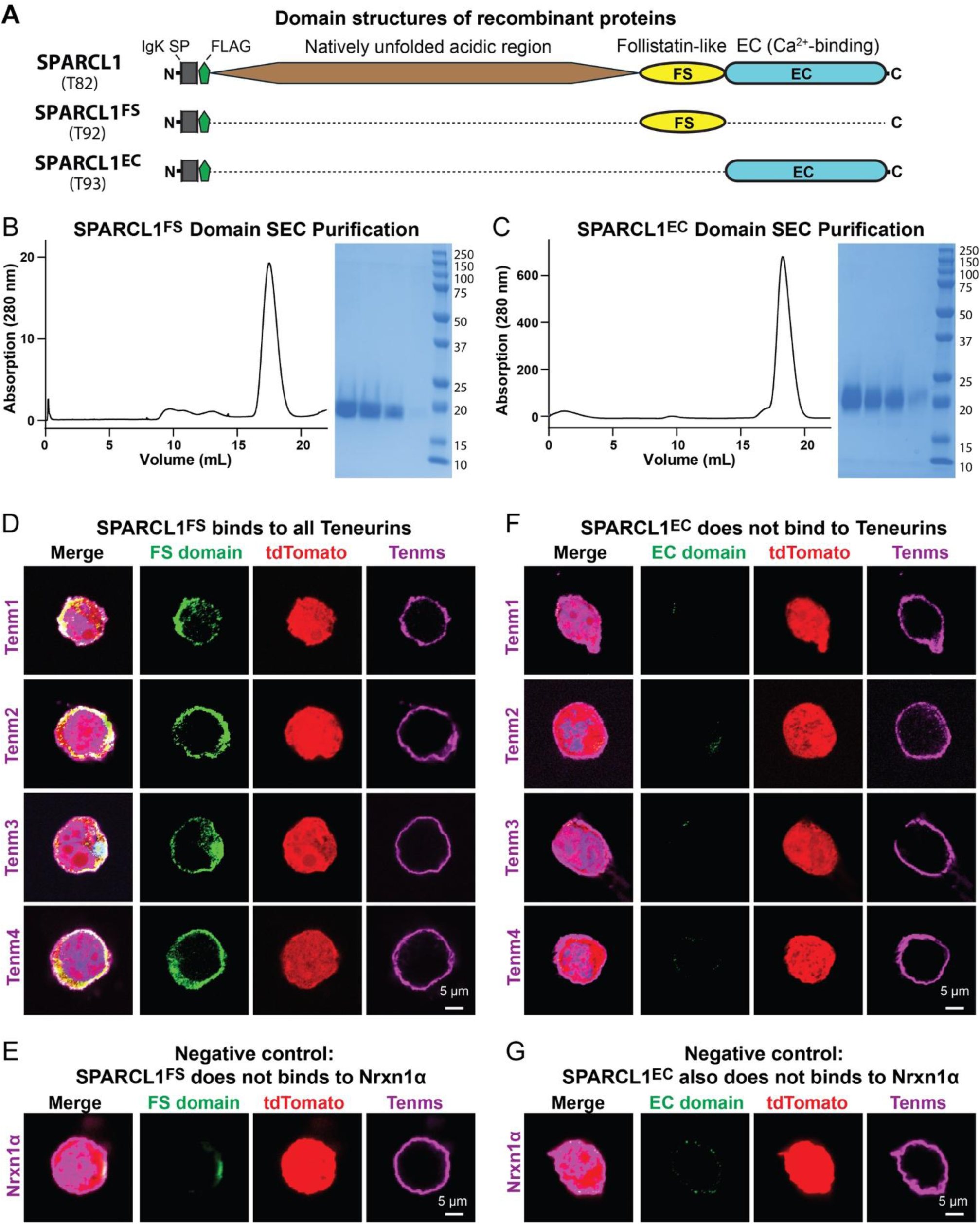
The follistatin-like (FS) domain of SPARCL1, but not its Ca^2+^-binding (EC) domain, avidly binds to all teneurins. (**A**) Schematic of full-length SPARCL1 and of follistatin-like (FS) and Ca^2+^-binding (EC) domain SPARCL1 proteins. All proteins were FLAG-tagged. (**B**,**C**) Purification of recombinant follistatin-like (FS) domain (B) and Ca^2+^-binding (EC) domain SPARCL1 proteins (C) by size-exclusion chromatography (SEC, left panels) and analysis of the purified proteins by Coomassie Blue-stained SDS-PAGE (right panels). (**D**-**F**) Cell-surface binding assays showing that the purified follistatin-like SPARCL1 domain protein binds robustly to all four teneurins (D) whereas the purified Ca^2+^-binding EC domain protein does not (F), and that neither protein binds to neurexin-1α (E, G). The assays were carried out as described for Figures 3 and 4 using 50 nM of recombinant proteins. Images correspond to 3 independently performed experiments with the same results.

Is the binding of the SPARCL1 FS domain to teneurins enough to stimulate synapse formation? We performed experiments again with mixed hippocampal neuron-glia cultures that were incubated with 50 nM of the recombinant FS domain or EC domain proteins as described above, using full-length SPARCL1 as a positive control (Fig. 7A, Fig. S3). Neither the FS nor the EC domain was able to stimulate synapse formation, whereas the positive SPARCL1 control did stimulate synapses (Fig. 7B-D). Thus, the EC domain appears to act as an effector domain for the FS domain of SPARCL1 that connects the EC domain to synaptic teneurins.

**Figure 7.**
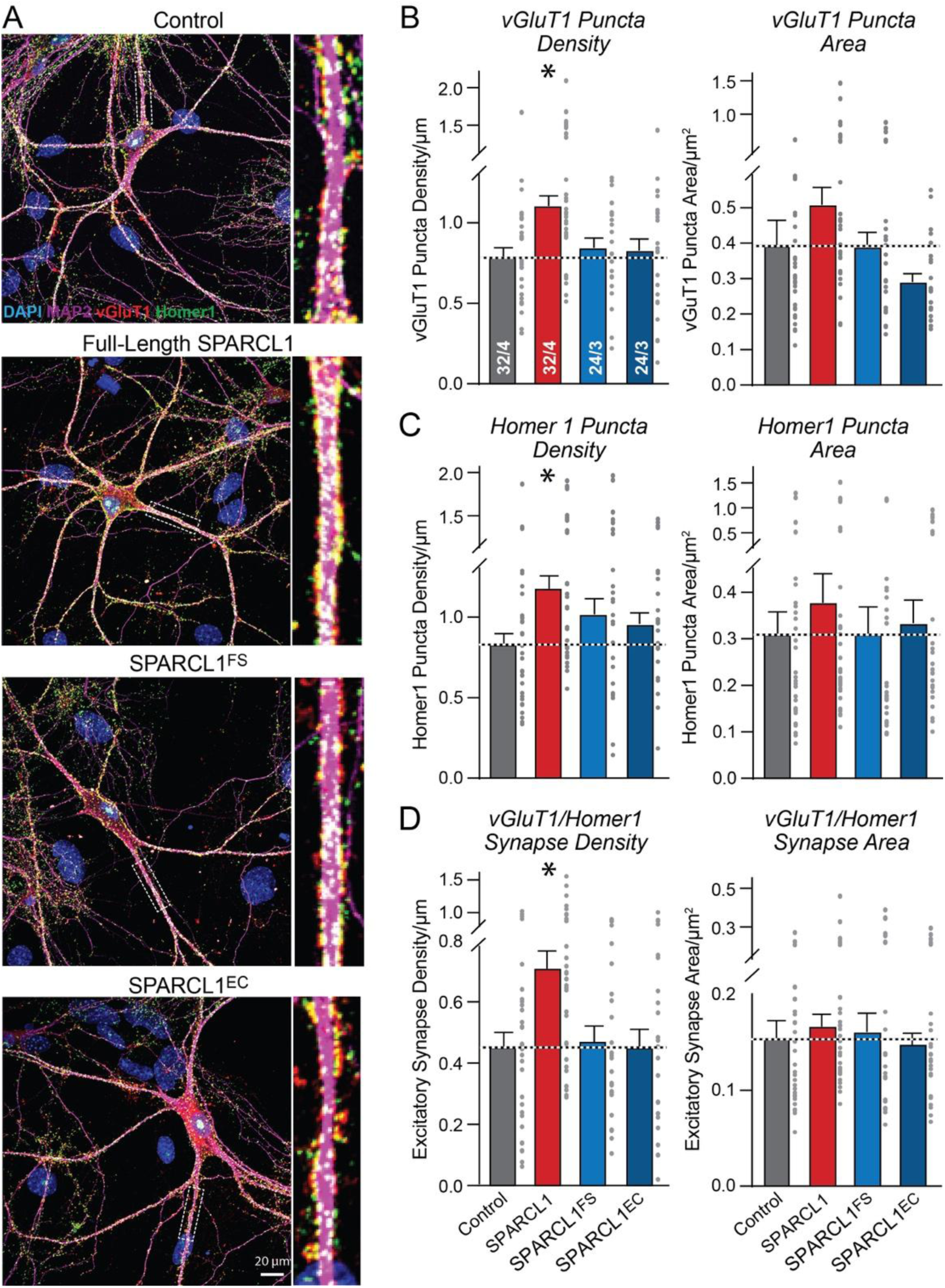
Teneurin-binding by the SPARCL1 follistatin-like domain is not sufficient for increasing synapse numbers. (**A**) Representative confocal images of mixed neuron-glia cultures treated with control medium or medium containing 50 nM recombinant full-length FLAG-tagged SPARCL1 or the isolated SPARCL1 follistatin-like domain (SPARCL1^FS^) or Ca^2+^-binding domain proteins (SPARCL1^EC^). Treatments of hippocampal cultures from newborn mice were performed and analyzed as described for Figure 1. For each condition, overviews are shown on the left and higher-magnification images taken from the boxed areas on the right. (**B-D**) Summary graphs of the synapse density and size of excitatory synapses show that full-length SPARCL1 but not the SPARCL1^FS^ or the SPARCL1^EC^ domain proteins robustly increase the excitatory synapse density and modestly increase the apparent synapse size (**B** & **C**, quantifications of the vGluT1- and Homer1-positive puncta, respectively; **D**, quantifications of synaptic puncta that are positive for both vGluT1 and Homer1, with all images recorded with identical parameters). Data are means ± SEM (numbers of cells and experiments are indicated in bars); *P < 0.05 [one-way ANOVA with post-hoc Tukey tests]. For single channel images, see Figure S3.

## DISCUSSION

The overall aim of the present study was to gain an understanding of how SPARCL1 (a.k.a. Hevin), a secreted protein primarily produced by astrocytes and vascular cells, induces synapse formation. These experiments were motivated by apparently incompatible results in the literature (see Introduction) and by the need to acquire insight into the powerful synaptogenic ability of SPARCL1 that dramatically boosts synapse numbers in cultured neurons (Fig. 1B). SPARCL1 consists of an N-terminal natively unfolded region that is poorly conserved and accounts for two thirds of its sequence and C-terminal FS and EC domains that are highly conserved and are similarly found in SPARC (after which SPARCL1 was named) and testicans (Fig. 8A). In this study, we made three overall observations that suggest a model for how SPARCL1 boosts synapse numbers:

**Figure 8.**
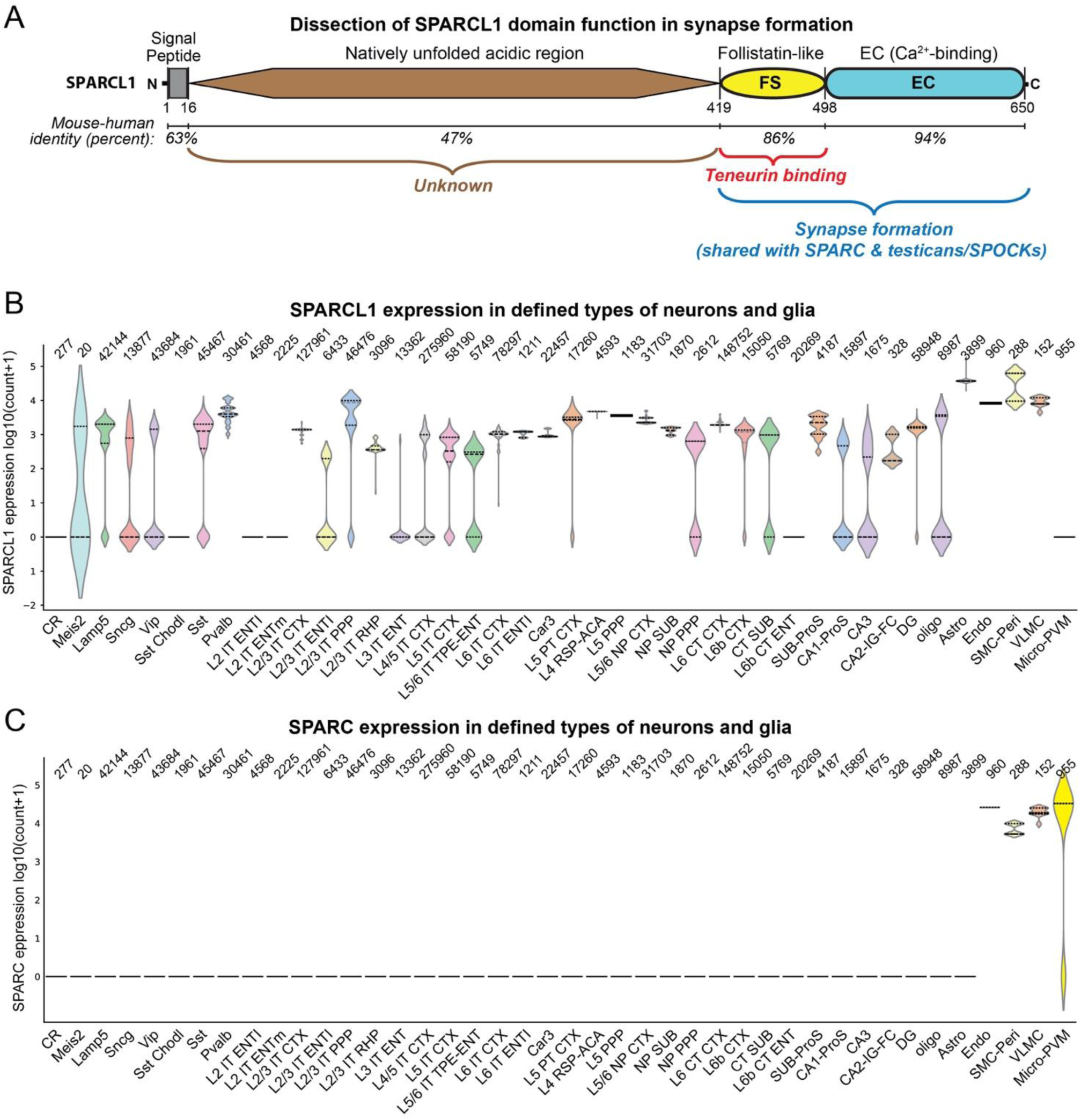
Model of SPARCL1 function in synapse formation and expression profile. (**A**) Summary model proposing that SPARCL1 follistatin-like (FS) domain binds to teneurins, while both SPARCL1 follistatin-like (FS) and Ca^2+^-binding (EC) domains are required for synapse formation. (**B**,**C**) Violin plots showing the expression levels of SPARCL1 (B) and SPARC (C) across major brain cell types, based on single-cell RNA-sequencing data from the Allen Brain Institute’s Whole Cortex & Hippocampus -10x Genomics (2020) dataset, annotated using the 10x–SMART-seq taxonomy (2021 release) (https://brain-map.org/our-research/cell-types-taxonomies/cell-types-database-rna-seq-data). The subclasses and cell-types are indicated in the supplementary table. The upper and lower edges mark the 75th and 25th percentiles; the number of cells is indicated above the violin plots. For assignment of the different cell types, please see the supplementary Excel file.

First, we find that the C-terminal FS and EC domains of SPARCL1 suffice to induce synapse formation at nanomolar concentrations (Fig. 1, 2). This result is consistent with the pattern of conservation of SPARCL1, although it does not quite agree with other papers Kucukdereli et al. (2011) and Singh et al. (2017) that localized the synaptogenic activity of SPARCL1 to a sequence N-terminal of the FS domain and found that the C-terminal domains are actually inhibitory for synapse formation. It is possible that these differences in findings are due to the differences in experimental preparations since we employed hippocampal cultures, whereas Kucukdereli et al. (2011) used cultured retinal ganglion cells.

Second, we find that at the synaptogenic concentration (50 nM), SPARCL1 binds to all teneurins but poorly interacts with neurexins or neuroligins, which were proposed as SPARCL1 receptors (Fan S,Gangwar SP,Machius M and Rudenko G, 2021) (Fig. 3-5). Since we also detected weak, possibly non-specific binding of SPARCL1 to Nrxn1α (Fig. 3A), it is possible that the differences in results here may be due to the concentrations used and that higher concentrations of SPARCL1 might promiscuously bind to multiple adhesion molecules. This hypothesis is consistent with the fact that the two papers reporting binding of SPARCL1 to neurexins and neuroligins (Fan S,Gangwar SP,Machius M and Rudenko G, 2021) localize the binding sites to distinct sequences of SPARCL1.

Third, we show that the small FS domain of SPARCL1 is sufficient for binding to teneurins but is not by itself capable of stimulating synapse formation. Instead, the FS domain requires coupling to the EC domain for synaptogenic activity (Fig. 6, 7). These observations suggest a model whereby recruitment of SPARCL1 to synapses via its FS domain-mediated interaction with teneurins enables the EC domain to induce or stabilize synapses.

Overall, our data confirm and extend Ben Barres’ pivotal observation that SPARCL1 is a powerful synaptogenic factor (Kucukdereli H et al., 2011). Reanalysis of recently available single-cell RNAseq data reveal that consistent with previous conclusions (Kucukdereli et al., 2011), SPARCL1 is highly expressed in astrocytes in brain but is also abundantly produced by endothelial and perivascular cells and by subclasses of neurons in brain (Fig. 8B), suggesting that SPARCL1 is not simply an astroglial synaptogenic factor but may be secreted by other types of cells as well, possibly in a regulated manner. SPARC, conversely, is only produced by perivascular cells and microglia, which in turn do not secrete SPARCL1 (Fig. 8C). The candidacy of teneurins as SPARCL1 receptors is attractive given that the deletion of teneurins produces extensive synapse loss (Zhang X et al., 2025;Zhang X et al., 2022), which is not the case for deletions of α-neurexins (Etherton MR et al., 2009;Missler M et al., 2003). Although innumerable papers have linked SPARCL1 to vascularization, inflammation, and cancer (Gagliardi F et al., 2017;Naschberger E et al., 2016;Zhao G et al., 2024), its potential receptors in these cells and its mechanism of action remain unknown. These cells do not express neurexins or neuroligins, but they do express teneurins and latrophilins, which are trans-cellular teneurin ligands (Arac D and Li J, 2019;Peppino G et al., 2021;Sudhof TC, 2025). It is thus possible that SPARCL1 also acts in non-neuronal cells by binding to teneurins, which would provide an attractive mechanism that explains the many functions of SPARCL1.

Our study, like many studies reporting unexpected results, raises new questions, two of which are particularly noteworthy. Most importantly, how does the EC domain of SPARCL1 induce synapses? The SPARCL1 EC domain is known to bind to collagen (Hambrock HO et al., 2003), raising the possibility of an involvement of the extracellular matrix in synapse formation. Another question regards the significance of the large N-terminal natively unfolded sequence of SPARCL1 which is lacking in SPARC. Does SPARCL1 have functions different from those of SPARC mediated by its large N-terminal sequence, or is this sequence functionally insignificant as suggested by its lack of evolutionary conservation?

Finally, our study has several limitations. Whereas we have shown that the deletion of neurexins or of neuroligins has no effect on the ability of SPARCL1 to induce synapse formation (Gan KJ and Sudhof TC, 2019;Gan KJ and Sudhof TC, 2020), we have not investigated whether the deletion of teneurins abolishes SPARCL1’s action because no conditional quadruple teneurin KO mice are available. Thus, deletion of all teneurin isoforms is technically difficult. Moreover, we have not tested the function of SPARCL1 in vivo. Cultures as a reduced system have the advantage that homeostatic mechanisms which adjust for overall synapse numbers and synaptic connectivity changes are not operating. As a result, many functions of molecules in synapses that are robust in cultured neurons as a reduced system are sometimes difficult to detect in vivo. Indeed, the reported phenotype of SPARCL1 knockout mice is modest, consistent with the supposition of compensatory mechanisms in vivo (Kucukdereli H et al., 2011;Ostrovskaya OI et al., 2020;Sullivan MM et al., 2008).

In sum, our findings extend previous results by outlining a potential mechanism of action of SPARCL1 in synapse formation but raise new questions that future studies will have to address.

## EXPERIMENTAL PROCEDURES

### Mouse Breeding and Husbandry

Male and female CD1 mice were weaned at 20-21 days of age and group-housed on a 12 h light-dark cycle with food and water ad libidum in the Stanford Veterinary Service Center. All procedures conformed to National Institutes of Health Guidelines for the Care and Use of Laboratory Mice and were approved by the Stanford University Administrative Panel on Laboratory Animal Care.

### Primary Mixed Neuron-Glia Cultures

The hippocampi of newborn male and female mice were dissected, digested with papain (Worthington) for 25 min at 37°C, and filtered through a 70 mm cell strainer (Falcon). Cells from male and female mice were mixed and plated in 24-well plates on coverslips that were coated with Matrigel (Corning). Plating media contained 5% fetal bovine serum (Sigma), B27 (Gibco), 0.4% glucose (Millipore-Sigma), and 2 mM glutamine (Gibco) in MEM (Gibco). One day after plating (DIV1), the culture medium was changed to growth medium containing B27 (Gibco), 2 mM glutamine (Gibco) in Neurobasal A (Gibco). At DIV4, half of the medium was exchanged by growth medium containing 4 mM Ara-C (Millipore-Sigma). Neurons were analyzed at DIV14-15. Hippocampal cells from both male and female newborn (P0) mice were combined to generate primary mixed neuron-glia cultures.

### Immunocytochemistry

All solutions were made fresh and filtered via a 0.22 µm filter before using for the experiments. Cells were washed once with DPBS, fixed with 4% PFA, 4% sucrose, and DPBS for 20 min at 4 °C, washed three times with DPBS, and permeabilized in 0.2% Triton X-100 in DPBS for 5 min at room temperature. Surface staining was performed in unpermeabilized samples (without 0.2% Triton X-100). Cells were subsequently placed in blocking buffer containing 4% goat serum (Millipore-Sigma), 3% BSA and DPBS for 1 h at room temperature, incubated with diluted primary antibodies (Rabbit anti-HA, Cell Signaling Technologies, #3724; Mouse anti-HA, Cell Signaling Technologies, #2367; Chicken anti- MAP2, EnCor Biotechnology Inc, #AB_2138173; Guinea pig anti-VGLUT1, Millipore, #AB5905; Rabbit anti-HOMER1, Millipore, #ABN37; Mouse anti-HOMER1, Synaptic Systems, #160011; Mouse anti-V5, Invitrogen, #R960CUS; Rabbit anti-ALFA, homemade; Mouse anti-FLAG, Sigma, #F3165; Rabbit anti-FLAG, Sigma, F7425) in blocking buffer overnight at 4 °C, washed three times with DPBS, incubated with diluted fluorescently-conjugated secondary antibodies (Goat anti-Chicken, Alexa Fluor™ Plus 488, Invitrogen, #A-32931; Goat anti-Guinea Pig, Alexa Fluor™ 546, Invitrogen, #A-11074; Goat anti-Rabbit, Alexa Fluor 647, Invitrogen, #A-21245; Goat anti-Mouse, Alexa Fluor™ 546, Invitrogen, #A- 11030; Goat anti-Rabbit, Alexa Fluor™ 546, Invitrogen, #A-11010; Goat anti-Guinea Pig, Alexa Fluor™ 647, Invitrogen, #A-21450; Goat anti-Mouse Alexa Fluor™ 647 Invitrogen, #A21236) in a blocking buffer for 2 h at room temperature, washed three times with DPBS, briefly dry and mounted on UltraClear microscope slides (Denville Scientific) using DAPI Fluoromount-G (Southern Biotech).

### Confocal Image Acquisition and Analysis

Serial confocal z-stack images were acquired using a Nikon confocal microscope (A1RSi) with a 60x oil-immersion objective. Images were analyzed using NIS-Elements AR acquisition software. Laser intensities and acquisition settings were established for individual channels using optimal LUT settings and applied to entire experimental replicates. All staining processes and acquisition parameters were kept constant among different experiments. For synaptic puncta quantification, z-stack images were acquired at 0.2 µm intervals, and ten consecutive sections with the highest signal were projected maximally to cover the strongest intensity signals. For HEK cell surface-binding assays, a single section of imaging was acquired. Image processing and quantification were performed using NIS-Elements and ImageJ with standardized filtering parameters.

### Calcium Imaging

Calcium imaging in hippocampal neurons was performed as described. Cultured neurons were infected with lentiviruses expressing hSyn-GCaMP6m at DIV4. Calcium imaging was performed at DIV14-15 using a Leica microscope at 37°C under 5% CO^2^. For potentiating synaptic transmission in cultured neurons, the cells were imaged at an ambient 4 mM CaCl_2_ and 8 mM KCl concentration equilibrated in a HEPES-based buffer (129 mM NaCl, 25 mM HEPES-NaOH, 15 mM glucose, 1 mM MgCl_2_, 10 μM glycine, adjust pH to 7.2 - 7.4) for 2 min. Image analysis was performed using MATLAB.

### Protein Expression and Purification

All secreted proteins were purified from conditioned culture medium using a standard His-tag affinity purification protocol adapted from Zhang et al. (2025). HEK293 cells were transfected at a density of 3.0×10^6^ cells/mL in a total culture volume of 50 mL. For each transfection, 270 µL of PEI MAX 40K (1 mg/mL) was mixed with 50 µg plasmid DNA in 5 mL Opti-MEM and incubated for 10 min at room temperature before addition to the cells. 18 hours after transfection, cultures were supplemented with 850 µL valproic acid (50 mg/mL) and 325 µL sodium propionate (100 mg/mL). Proteins were expressed for 96 h at 37 °C following transfection. For purification, conditioned media were clarified by centrifugation and adjusted to pH 8.3 using 50 mM Tris-HCl. The supernatants were incubated overnight at 4 °C with gentle agitation in the presence of Ni^2+^-NTA resin (Qiagen, #30210). Resin-bound proteins were collected using gravity-flow columns, washed with wash buffer containing 50 mM Tris-HCl (pH 8.3), 500 mM NaCl, and 8 mM imidazole, and eluted with elution buffer containing 50 mM Tris-HCl (pH 8.3), 500 mM NaCl, and 500 mM imidazole. Eluted proteins were dialyzed into general protein buffer containing 50 mM HEPES-NaOH (pH 7.3) and 100 mM NaCl, concentrated using centrifugal concentrators, and further purified by size-exclusion chromatography (SEC) using an NGC chromatography system (Bio-Rad) equipped with a Superdex 200 Increase column (Cytiva) equilibrated in the same buffer.

### Data Analyses and Statistics

All bar plots show means ± SEM except the single-cell RNA analyses in Fig. 8. For all violin plots, the central line shows the median and the upper and lower lines show the quartiles (25th and 75th percentiles). Statistical analyses were performed with GraphPad Prism 10 software using unpaired two-tailed Student’s t-tests for experiments with only two groupsand one-way ANOVA with Tukey’s multiple comparison tests to compare the mean of each column with the mean of every other column for experiments with more than two groups. All numbers of cells, neurons, and replicates analyzed are shown in the bars or plots. All experiments were performed and analyzed in a blind manner by an experimenter except for the immunoblots for which this is not possible. All figures and data were scanned by ‘Proofig’ software to avoid accidental copy-paste errors in the assembly of figures, tables, and graphs.

## Data Availability

All raw data for this study are publicly available at …. (Stanford Digital Repository - to be filled in upon publication of the study).

## ACKNOWLEDGEMENTS

We would like to thank Dr. Kathlyn Gan for her initial contributions to this project. This work was supported by a grant from the NIMH (5R01 MH126929-02 to T.C.S.) and a fellowship from the Larry L. Hillblom Foundation (2022-A-015-FEL to X.Z.).

## AUTHOR CONTRIBUTIONS

Conceptualization: X.Z., and T.C.S.; Methodology: X.Z., X.C., Y.M., and T.C.S.; Investigation: X.Z., X.C., and Y.M.; Funding acquisition: T.C.S.; Supervision: X.Z. and T.C.S.; Writing - review and editing: X.Z., X.C., Y.M., and T.C.S.

## CONFLICT OF INTEREST

The authors declare that they have no competing interests.

---

**Figure S1.**
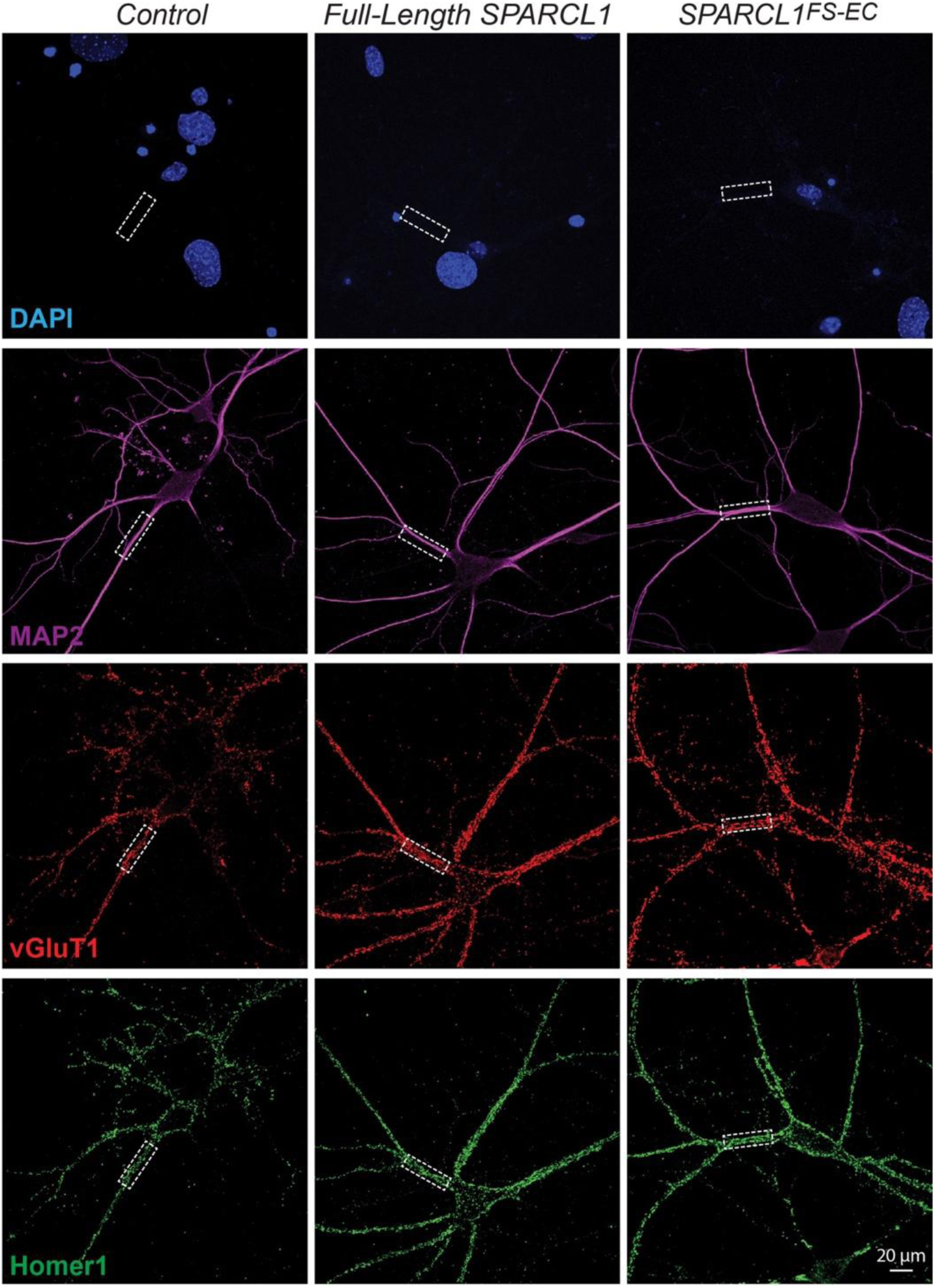
Single channel images for Figure 1. Images depict the individual single channels for the data shown in Figure 1B. Neurons were stained for DAPI (blue), MAP2(Magenta), vGluT1 (red), and Homer1 (green). Left, with control medium; middle, with 50 nM full-length SPARCL1 protein; right, with 50 nM C-terminal SPARCL1^FS-EC^ protein.

**Figure S2.**
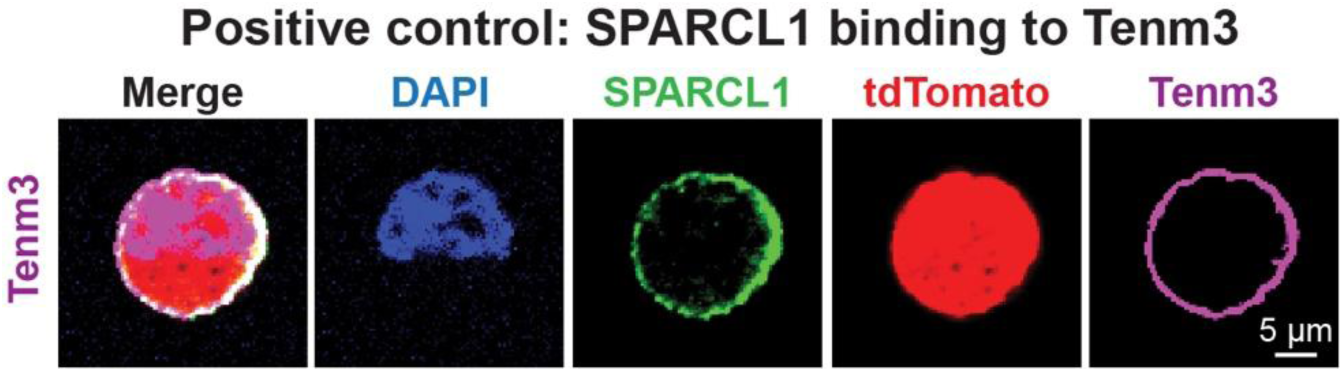
Additional positive control for Figure 3. Representative images from experiments independently repeated 3 times showing that teneurin-3 (Tenm3) binds to SPARCL1.

**Figure S3.**
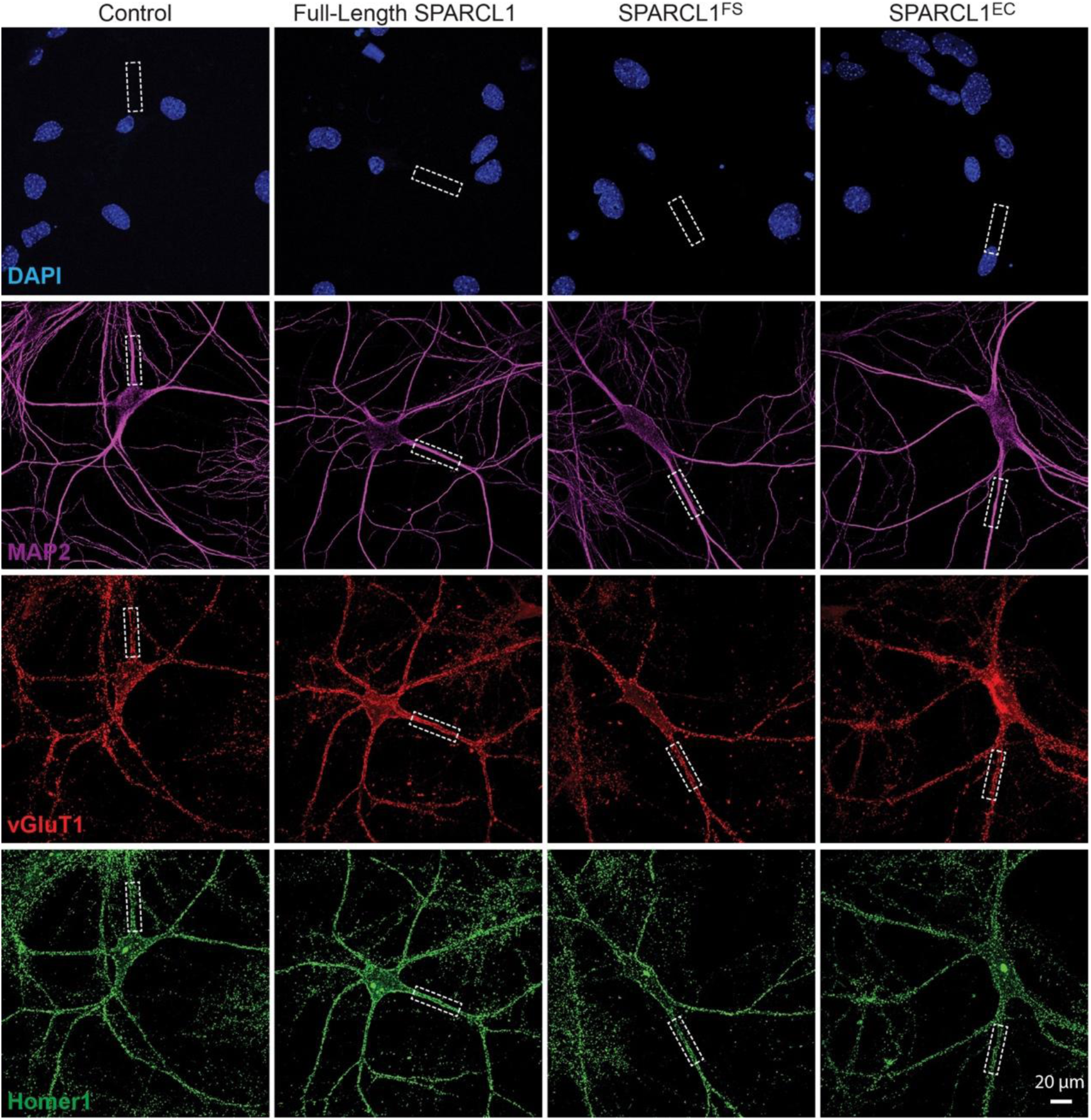
Single channel images for Figure 7. Images depict the individual single channels for the data shown in Figure 7A. Neurons were stained for DAPI (blue), MAP2 (magenta), vGluT1 (red), and Homer1 (green). Left, with control medium; middle, with 50 nM full-length SPARCL1 protein; middle-left, with 50 nM full-length SPARCL1 protein; middle-right, with 50 nM SPARCL1^FS^; right, with 50 nM SPARCL1^EC^ protein.

## Notes

### Competing Interest Statement

The authors have declared no competing interest.

